# Involvement of uterine natural killer cells in the natural selection of human embryos at implantation

**DOI:** 10.1101/2020.08.14.251033

**Authors:** Chow-Seng Kong, Alexandra Almansa Ordoñez, Sarah Turner, Tina Tremaine, Joanne Muter, Emma S. Lucas, Emma Salisbury, Rita Vassena, Ali A. Fouladi-Nashta, Gustavo Tiscornia, Geraldine Hartshorne, Jan J. Brosens, Paul J. Brighton

## Abstract

Decidualizing endometrial stromal cells (EnSC) critically determine the maternal response to an implanting conceptus, triggering either menstruation-like disposal of low-fitness embryos or creating an environment that promotes further development. However, the mechanism that couples maternal recognition of low-quality embryos to tissue breakdown remains poorly understood. Recently, we demonstrated that successful transition of the cycling endometrium to a pregnancy state requires selective elimination of pro-inflammatory senescent decidual cells by activated uterine natural killer (uNK) cells. Here we report that uNK cells express CD44, the canonical hyaluronan (HA) receptor, and demonstrate that high-molecular weight HA (HMWHA) inhibits uNK cell-mediated killing of senescent decidual cells. By contrast, low-molecular weight HA (LMWHA) did not attenuate uNK cell activity in co-culture experiments. Killing of senescent decidual cells by uNK cells was also inhibited upon exposure to medium conditioned by IVF embryos that failed to implant, but not successful embryos. Embryo-mediated inhibition of uNK cell activity was reversed by recombinant hyaluronidase 2 (HYAL2), which hydrolyses HMWHA. We further report a correlation between the levels of HYAL2 secretion by human blastocysts, morphological scores, and implantation potential. Taken together, the data suggest a pivotal role for uNK cells in embryo biosensing and endometrial fate decisions at implantation.

## Introduction

The midluteal window of implantation constitutes an inflection point in the menstrual cycle, after which the endometrium either breaks down or becomes transformed into a semipermanent tissue, i.e. the decidua of pregnancy.^1^ The fate of the endometrium depends foremost on implantation of an embryo capable of producing sufficient human chorionic gonadotrophin (hCG), and other fitness hormones, to rescue the corpus luteum and maintain ovarian progesterone production.^2^ Although progesterone exerts multiple functions, its primary role is to drive differentiation of endometrial stromal cells (EnSC) into specialised decidual cells,^1^ which first encapsulate the implanting blastocyst^3,4^, and then form a robust immune-protective matrix to accommodate invading placental trophoblast.^1,5^

Recent studies demonstrated that decidualization is a multistep differentiation process that starts with an evolutionarily conserved acute cellular stress response,^6^ exemplified by a burst of free radicals and release of inflammatory mediators.^7,8^ This inflammatory phase coincides with the implantation window,^9,10^ and is characterised further by accumulation of proliferating uterine natural killer (uNK) cells.^8^ After a lag-period of approximately four days, phenotypic decidual cells emerge that are progesterone-dependent,^11^ resistant to oxidative and metabolic stress,^12,13^ and highly responsive to embryonic signals.^3,4,14^ However, inflammatory reprogramming of EnSC also leads to the emergence of acute senescent cells.^8,15^ The topography of senescent decidual cells, i.e. in proximity to the luminal epithelium, suggests that they arise from cells subjected to significant replication stress during the proliferative phase.^15^ Although morphologically similar to decidual cells, senescent decidual cells are progesterone-resistant and secrete an abundance of extracellular matrix proteins and proteinases, proinflammatory cytokines, and chemokines (designated senescence associated secretory phenotype or SASP), which cause sterile inflammation and induce secondary senescence in neighbouring decidual cells.^8,15,16^ Under continuous progesterone signalling in pregnancy, decidual cells secrete IL-15 and other factors involved in recruitment and activation of uNK cells, which in turn target and eliminate senescent decidual cells through perforin- and granzyme-containing granule exocytosis.^8,15^ In a non-conception cycle, however, falling progesterone in the late-luteal phase leads to a preponderance of senescent cells, influx of leukocytes, extracellular matrix (ECM) breakdown, and menstrual shedding of the superficial uterine mucosa.^17^ These novel insights into the cellular dynamics in the stroma during the window of implantation place uNK cells at the centre of endometrial fate decisions.

Human preimplantation embryos are genetically diverse and prompt elimination of embryos of low-fitness is paramount to limit maternal investment in failing pregnancies.^18,19^ Several studies have highlighted that decidual cells are programmed to respond to embryos of different quality.^3,4,14,20,21^ Further, growing evidence indicates that loss of this biosensing function of the endometrium causes recurrent pregnancy loss.^18,22^ Whether uNK cells are involved in embryo selection at implantation is not known, although there is evidence that hCG stimulates uNK cell proliferation.^23^

Hyaluronan (HA), a ubiquitous ECM component, plays an important role in diverse biological processes that involve rapid tissue turnover and repair.^24^ Not surprisingly, HA has been implicated in key reproductive events, including ovulation, fertilization, embryogenesis, implantation, and trophoblast invasion.^25–27^ HA, a linear glycosaminoglycan consisting of repeating disaccharide units of D-glucuronic acid and N-acetyl-D-glucosamine, is expressed in various molecular sizes up to 10^7^ kDa.^25^ It is produced by three different hyaluronic acid-synthase enzymes (HAS1-3), each generating polymers of different lengths. Turnover of HA at tissue level is rapid, mediated by a family of hyaluronidases (HYALs). The biological actions of HA depend on its size and concentration.^24,25^ High-molecular weight HA (HMWHA) polymers are space-filling hydrating molecules that impede cell differentiation and exert anti-angiogenic and immunosuppressive activities. On the other hand, low-molecular weight HA (LMWHA; molecular mass ≤ 250 kDa) activates a number of receptors, most prominently the cell-surface glycoprotein CD44 and hyaluronan mediated mobility receptor (HMMR; otherwise known as RHAMM or CD168), involved in cell proliferation and migration, angiogenesis, and immunomodulation.^24,25^

In this study, we report that downregulation of HAS2 expression upon decidualization of EnSC leads to loss of HA production. We exploited this observation to demonstrate that exogenous HMWHA, but not LMWHA, abrogates uNK cell-mediated killing of senescent decidual cells. Further, we show that human IVF blastocysts of poorer morphological quality and those that failed to implant secrete lower levels of hyaluronidase 2 (HYAL2) when compared to their successful counterparts. Importantly, the secretome of unsuccessful embryos also strongly inhibited uNK cell activity, an effect reversed by recombinant HYAL2. By contrast, we found no evidence that successful embryos impede uNK cell-mediated clearance of senescence decidual cells.

## Results

### Loss of HA synthesis upon decidualization of EnSC

We examined HA expression in decidualizing primary EnSC cultures using biotin-labelled hyaluronic acid binding protein (HABP).^28^ HABP reactivity was prominent in undifferentiated EnSC but downregulated upon decidualization with 8-bromo-cAMP and the progestin medroxyprogesterone (C+M) (Fig. 1A). Quantitative analysis confirmed that HA reactivity declines by 74% and 94%, respectively, after 4 and 8 days of decidualization (Fig. 1B). To explore the mechanisms accounting for the loss of HA production upon decidualization, we examined the expression of the three hyaluronic synthases (HAS1-3) and five catabolic hyaluronidases [HYAL1-4 and sperm adhesion molecule 1 (SPAM1)].^25,29,30^ First, we mined publicly available RNA-seq data of undifferentiated EnSC and cells decidualized for 4 days (GEO accession number: GSE104721). *HAS1, HAS3, HYAL1* and *HYAL4* mRNA levels were generally below 3 transcripts per million (TPM) in both undifferentiated and decidualized cells, suggesting little or no expression (Fig. S1). *SPAM1*, a testis-specific gene,^31^ is not expressed by EnSC and *HYAL3* does not encode for a classical hyaluronidase, although it may play a role in augmenting HYAL1 activity.^32,33^ By contrast, the average (± SEM) expression of *HAS2* and *HYAL2* in undifferentiated EnSC was 20.5 (±11.4) and 48.0 (±4.3) TPM, respectively. Upon decidualization, *HAS2* was downregulated whereas *HYAL2* transcript levels remained unchanged (Fig. S1). To confirm these observations, *HAS2* and *HYAL2* mRNA levels were measured by RT-qPCR in five independent primary EnSC cultures decidualized for 4 or 8 days. In agreement with the RNA-seq data, *HYAL2* expression was maintained throughout the decidual time-course whereas *HAS2* mRNA levels declined by 56% and 82% after 4 and 8 days of decidualization, respectively (Fig. 1C; *P* < 0.005, Dunnett’s multiple comparisons test).

**Figure 1.**
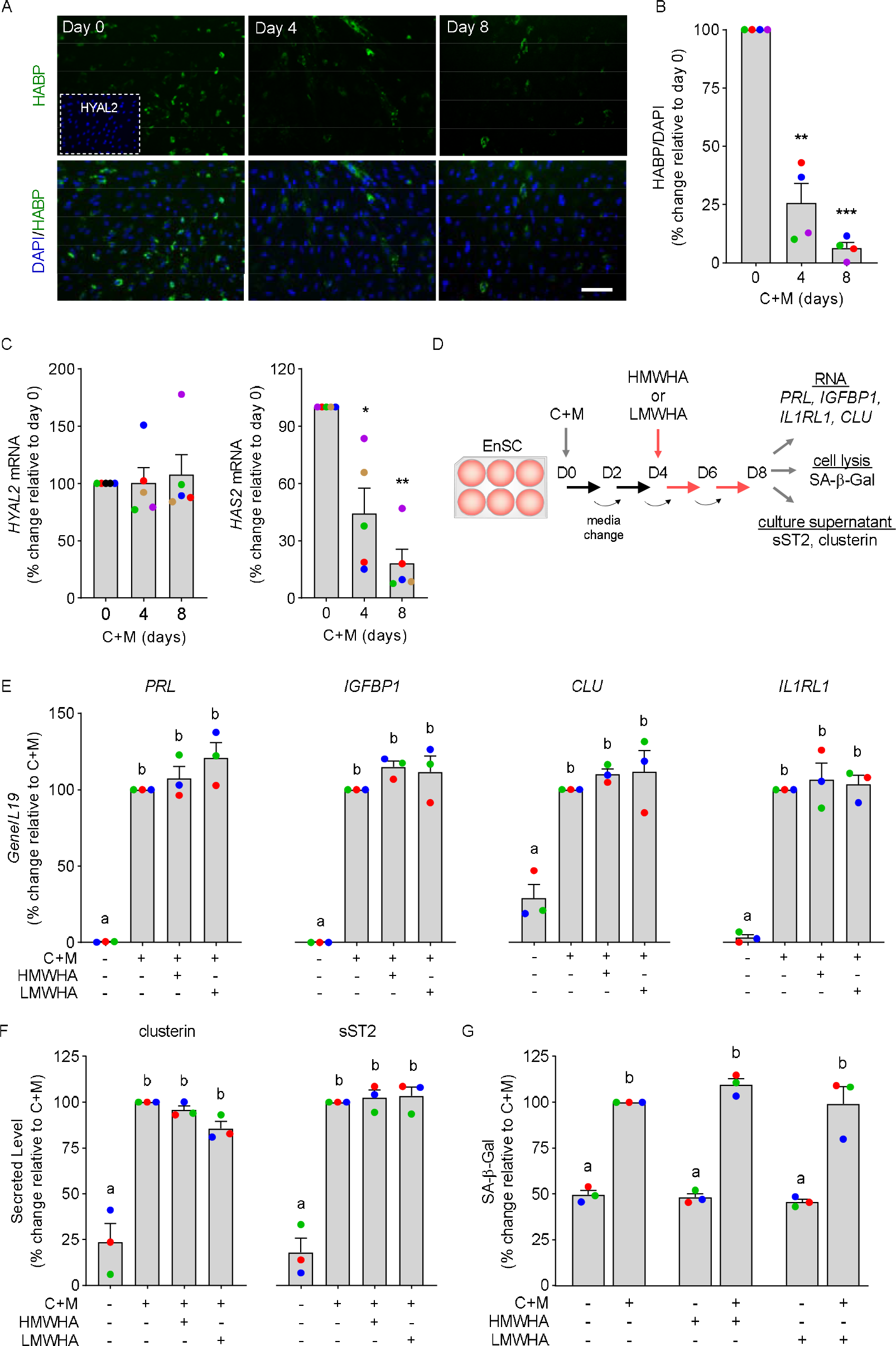
Loss of HA production upon decidualization. (**A**) Representative images of HABP immunofluorescence in undifferentiated EnSC (day 0) and cells decidualized with 8-br-cAMP and MPA (C+M) for 4 or 8 days. Scale = 200 μM. Cultures treated with recombinant HYAL2 to digest HA were used as control (*insert*). (**B**) Quantification of HABP^+^ cells. The data show the relative change in HABP^+^ cells (mean ± SEM) upon decidualization in 4 independent primary cultures. **indicates *P* < 0.01 and *** *P* < 0.001, ANOVA with Dunnett’s multiple comparison test. (**C**) RT-qPCR analysis of *HYAL2* mRNA (left panel) and *HAS2* mRNA (right panel) expression in undifferentiated EnSC (day 0) and cells decidualized for 4 or 8 days. The data show the relative change in transcript levels (mean ± SEM) upon decidualization of 5 independent primary cultures at the indicated timepoints. *denotes *P* < 0.05 and ** *P* < 0.01, ANOVA and Dunnett’s multiple comparison test. (**D**) Experimental design of decidualizing EnSC treated with HA. (**E-G**) HMWHA and LMWHA do not impact decidual marker gene expression (**E**), clusterin and sST2 secretion (**F**), or SA-β-Gal activity (**G**). Individual data points from 3 biological repeat experiments are shown together with bar graphs denoting mean ± SEM. Different letters indicate significance between groups at *P* < 0.05 (one-way ANOVA and Dunnett’s multiple comparison test).

We postulated that loss of endogenous HA production may render decidualizing cells responsive to HA derived from other sources, for example the implanting blastocyst.^34^ To test this hypothesis, three independent primary EnSC cultures were first decidualized for 4 days to downregulate endogenous HA production and then decidualized for 4 additional days in the presence or absence of LMWHA or HMWHA (Fig. 1D). As shown in Figure 1E, HA had no effect on the induction of the canonical decidual marker genes *PRL* and *IGFBP1*, irrespective of its molecular weight. Expression of *CLU* and *IL1RL1*, marker genes of senescent decidual cells and decidual cells, respectively,^15^ were also not affected by HA. *CLU* codes the secreted molecular chaperone clusterin (also known as apolipoprotein J) whereas both the transmembrane and soluble IL-33 receptors (termed ST2L and sST2, respectively) are encoded by *IL1RL1*. In keeping with the observations at transcript level, we found no evidence that HA impacted on either clusterin or sST2 secretion in decidualizing cells (Fig. 1F). To confirm that HA does not affect decidual subpopulations, senescence-associated β-galactosidase (SA-β-Gal) activity was measured in parallel cultures. As expected, decidualization was associated with a robust increase in SA-β-Gal activity but addition of HA had no effect on basal or induced levels (Fig. 1G).

### HMWHA inhibits uNK cell clearance of senescent decidual cells

To explore if there is a role for HA signalling in the peri-implantation endometrium, we examined the expression of putative HA receptors using recently generated single-cell RNA-seq data from endometrial biopsies obtained 8 and 10 days following the pre-ovulation LH surge (GEO accession number: GSE127918).^15^ Several genes coding HA receptors were expressed prominently in endometrial endothelial cells (Fig. 2A), including *ICAM1* (coding intercellular adhesion molecule 1), *LAYN* (layinin) and *TLR4* (toll-like receptor 4), pointing towards a potential role for HA signalling in vascular remodelling in pregnancy. The most striking observation, however, was the high expression of CD44, the principal HA receptor, in uNK cells (Fig. 2A). CD44 regulates cytotoxicity of circulating NK cells and potentially their trafficking to the endometrium.^35,36^ These observations prompted us to investigate if HA-CD44 interactions modulate uNK cell-dependent clearance of senescent decidual cells. To test our hypothesis, uNK cells were isolated from midluteal endometrial biopsies using magnetic-activated cell sorting (MACS). Flow cytometry analysis of uNK cells isolated from 3 independent biopsies showed an average viability (± SEM) of 80.8% (± 2.5) (Fig. 2B). Further, 95% (± 1.4) of CD56^+^ cells co-expressed the pan-leukocyte marker CD45 (Fig. 2C) and 79.7% (± 1.9) the canonical HA receptor CD44 (Fig. 2D). Expression of CD44 on CD56^+^ uNK cells was confirmed by immunocytochemistry (Fig. 2E). To validate that uNK cells selectively target senescent decidual cells, co-cultures were established with EnSC first decidualized for 6 days. After 48 h, the co-cultures were processed for SA-β-Gal activity measurement or RNA extraction (Fig. 2F). Spent medium was also collected and secreted clusterin and sST2 levels measured by ELISA. As expected, the induction of SA-β-Gal activity upon decidualization was abrogated in the presence of uNK cells, as was the induction and secretion of clusterin (Fig. 2H). By contrast, *IL1RL1* mRNA expression and sST2 levels were unaffected (Fig. 2I), confirming that uNK cells selectively target and eliminate senescent decidual cells.

**Figure 2.**
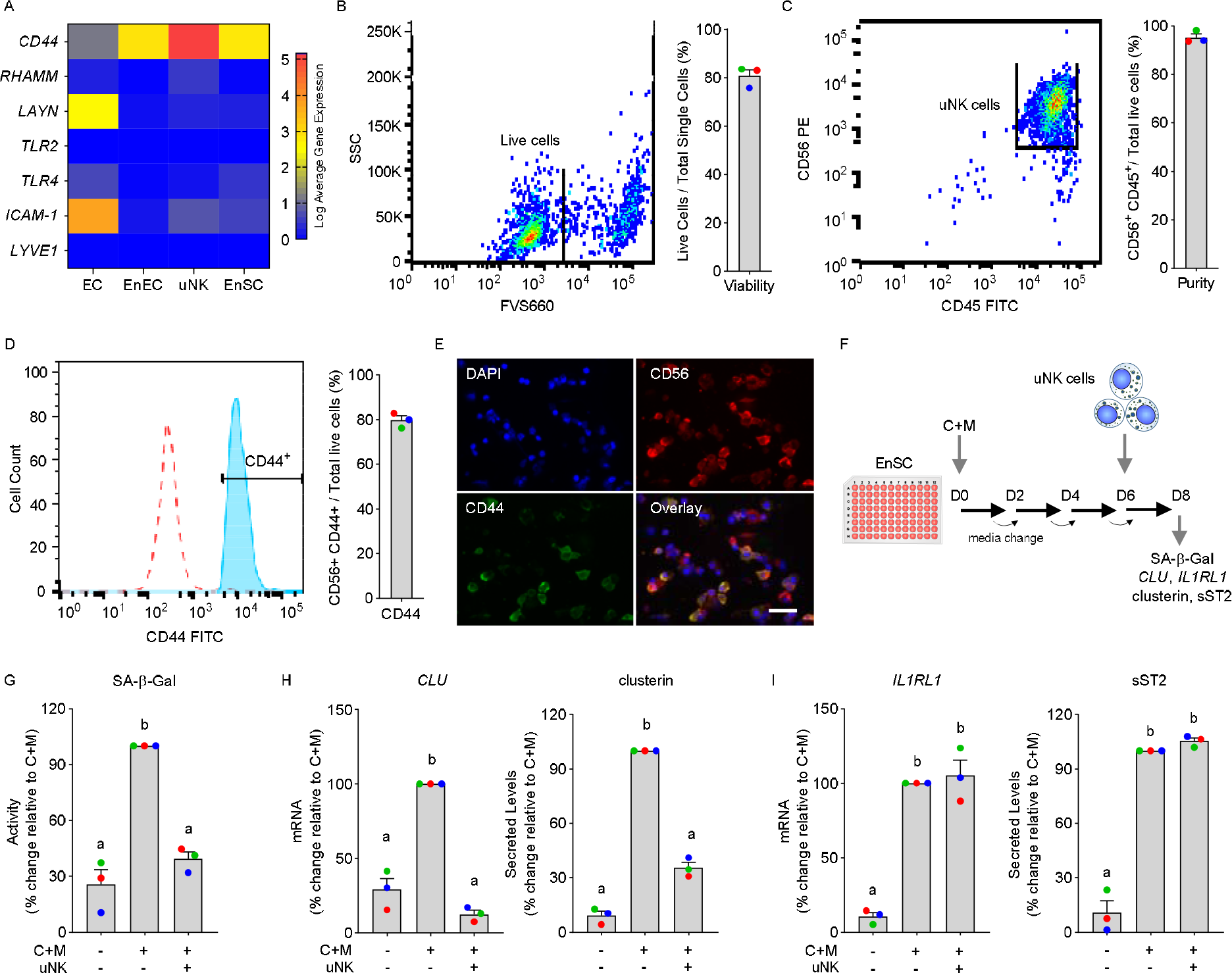
Decidual-uNK cell co-cultures. (**A**) Heatmap showing relative expression of genes coding putative HA receptors in different endometrial cell types *in vivo*. EC, endothelial cells; EnEC, endometrial epithelial cells. (**B-D**) Flow cytometric analysis. (**B**) Viability of uNK cells following MACS isolation was determined by FV660 exclusion. (**C**) Purity of uNK cells after MACS isolation was determined by gating cells for CD56 and CD45. (**D**) Histogram depicting CD56^+^/CD44^+^ uNK cells after MACS isolation (*filled histogram*) versus negative control (*open histogram*). Representative dot plots or histogram are shown together with bar graphs (mean ± SEM) showing individual data points of 3 biological repeat experiments. (**E)** Cytospin preparations of isolated uNK cells stained for DAPI (nucleus), CD56 and CD44. Co-localization of CD56 and CD44 is shown in bottom right panel. Scale = 200 μM. **(F)** Schematic representation of the decidual-uNK cell co-culture killing assay. **(G-I)** Analysis of SA-β-Gal activity (**G**), senescent decidual cell markers (**H**), and decidual markers (**I**) in decidualizing EnSC co-cultured with or without uNK cells. The data show relative change to the indicated measurements in 3 biological repeat experiments compared to cultures decidualized with C+M for 8 days in the absence of uNK cells. Different letters above the bar graphs (mean ±SEM) indicate statistical differences (*P* < 0.05) between groups (one-way ANOVA and Dunnett’s multiple comparison test).

Next, we treated 7 independent decidual-uNK cell co-cultures with either HMWHA or LMWHA. As shown in Figure 3A, HMWHA inhibited uNK cell-mediated clearance of senescent decidual cells, exemplified by the lack of downregulation of SA-β-Gal activity and clusterin secretion. sST2 secretion was unaffected by HMWHA. Further, addition of LMWHA had no effect on either SA-β-Gal activity or clusterin and sST2 secretion in decidual-uNK cell co-cultures. To substantiate these observations, 6 additional co-cultures were established and treated with either HMWHA or HMWHA pre-incubated with recombinant HYLA2, the enzyme that selectively degrades HMWHA into smaller fragments.^37^ As demonstrated in Figure 3B, HMWHA inhibition of uNK cell-dependent clearance of senescent decidual cell markers, monitored by SA-β-Gal activity and clusterin secretion, was completely lost upon pre-incubation with recombinant HYLA2. Addition of HYAL2 alone to the co-cultures had no effect on uNK-dependent clearance of senescent decidual cells (Fig. 3B). Degradation of HA by HYAL2 into smaller fragments initiates CD44 cell signalling.^24,25,38^ To explore the role of CD44, co-cultures were treated with Hermes-1, a monoclonal antibody that recognizes the N-terminal HA binding domain of CD44.^39^ Incubation of co-cultures with this blocking antibody inhibited uNK cell-mediated clearance of senescent decidual cells (Fig. 3C), irrespective of treatment with LMWHA or HMWHA. Taken together, the data indicate that HMWHA is a potent inhibitor of uNK cell activation, an effect that likely involves silencing of CD44 signalling.

**Figure 3.**
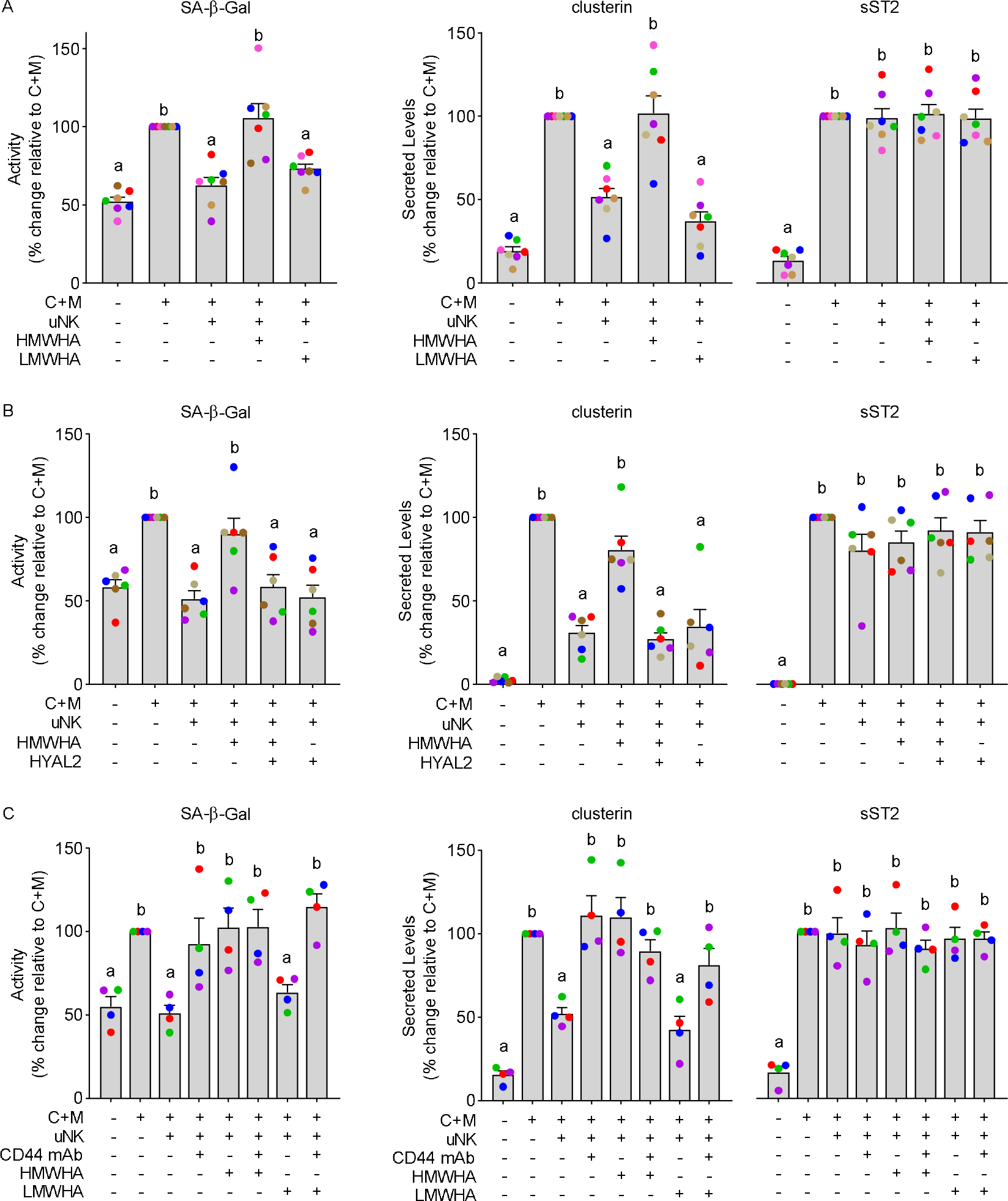
HMWHA inhibits uNK cell-dependent killing of senescent decidual cells. (**A**) Impact of HMWHA and LMWHA on uNK cell-dependent killing of senescent decidual cells in 7 biological repeat experiments. Killing of senescent decidual cells was monitored by SA-β-Gal activity and secreted clusterin levels. sST2 is selectively secreted by decidual cells. (**B**) Impact of HMWHA or HMWHA pre-incubated with HYAL2 on uNK cell-dependent killing of senescent decidual cells in 6 biological repeat experiments. (**C**) Impact of CD44 inhibition using a monoclonal antibody (CD44 mAb) on uNK cell-dependent killing of senescent decidual cells in the presence or absence of HMWHA or LMWHA in 4 biological repeat experiments. The data show relative change in the indicated measurements compared to cultures decidualized for 8 days in the absence of uNK cells. Different letters above the bar graphs (mean ± SEM) indicate statistical differences (*P* < 0.05) between groups (one-way ANOVA and Dunnett’s multiple comparison test).

### uNK cells are biosensors of embryo quality at implantation

The HA system is tightly controlled in pre-implantation embryos and trophoblast cells.^25,27^ We speculated that embryo-derived HA signalling may determine endometrial responses at the implantation site by modulating uNK cell activity. To investigate this possibility, we adapted our co-culture system as depicted in Figure 4A. Briefly, in two different IVF centres, blastocyst conditioned medium (BCM) was pooled from individual fully anonymised day-5 IVF embryos, which resulted either in an ongoing pregnancy following single-embryo transfer (BCM^+^, i.e. positive pregnancy test 15 days after embryo transfer; n = 28) or failed implantation (BCM^−^, negative pregnancy test; n = 38). Each pool consisted of 80 μL of spent media recovered from 7 blastocysts. We also collected unconditioned media (UCM) from parallel embryo-free droplets. Next, we decidualized 9 independent primary EnSC cultures for 6 days and then co-cultured uNK cells in UCM, BCM^+^ or BCM^−^ diluted 1 in 7.5 with decidualization medium. Co-cultures were harvested after 48 h. Strikingly, uNK cell-mediated killing of senescent decidual cells, as measured by SA-β-Gal activity and secreted clusterin levels, was unaffected by UCM or BCM^+^ but markedly inhibited in response to soluble signals emanating from unsuccessful blastocysts (BCM^−^) (Fig. 4B). Pre-incubation of BCM^−^ with recombinant HYLA2 fully restored uNK cell activity against senescent decidual cells (Fig. 4C). Notably, exposure of decidualizing EnSC to either UCM, BCM^+^ or BCM^−^ had no effect on either SA-β-Gal activity or clusterin and sST2 secretion in the absence of cocultured uNK cells (Fig. S2). Taken together, the data strongly infer a central role for uNK cells in embryo recognition and selection upon implantation. Further, the level of embryo-derived HYAL2 secretion appears to determine uNK cell activity.

**Figure 4.**
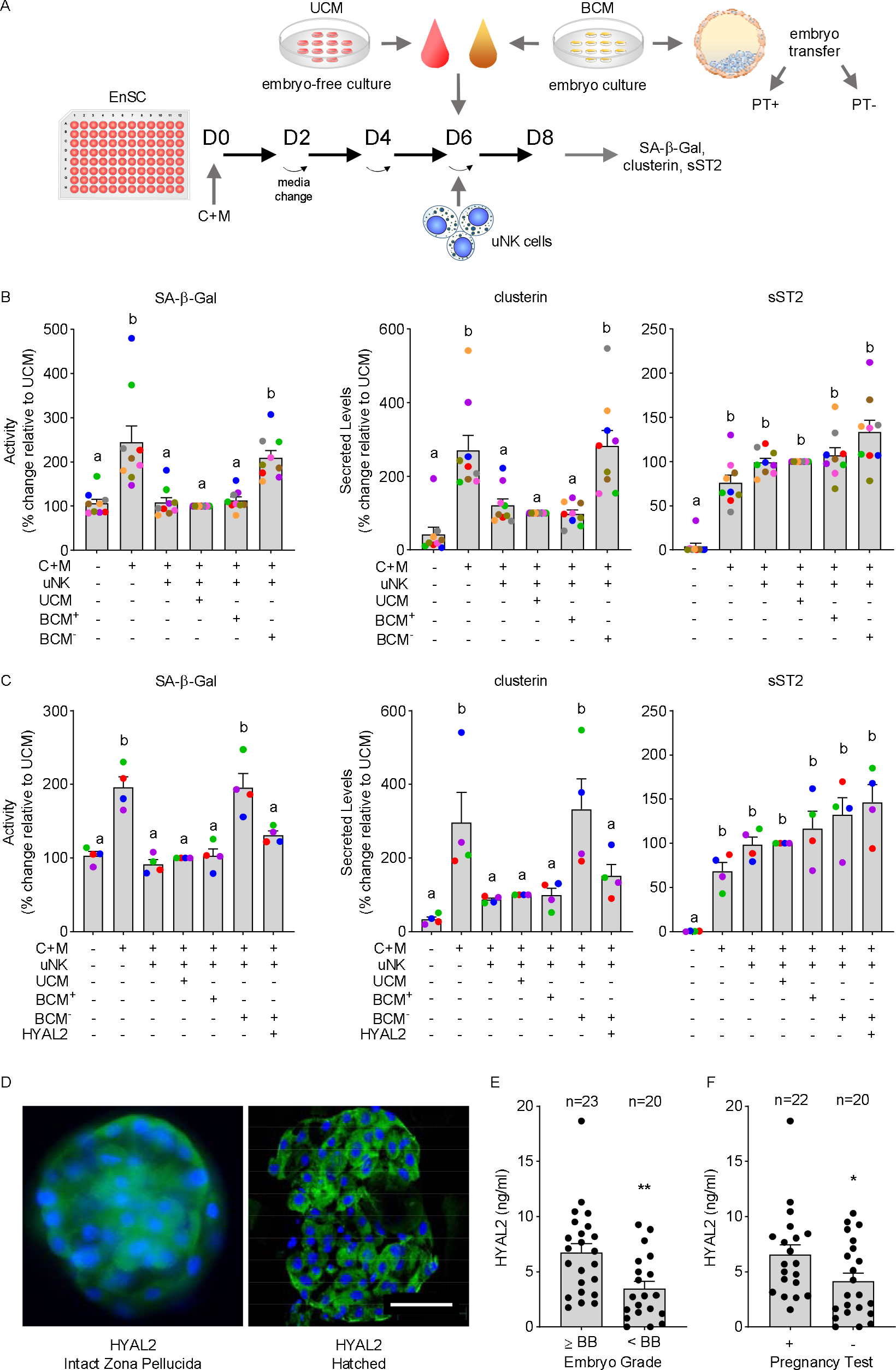
uNK cells are biosensors of low-fitness human embryos. (**A**) Schematic depiction of experimental design. BCM, blastocyst conditioned medium; UCM, unconditioned medium; PT, pregnancy test. (**B**) Impact of UCM, BCM^+^, and BCM^−^ on uNK cell-dependent killing of senescent decidual cells in 9 biological repeat experiments. Killing of senescent decidual cells was monitored by SA-β-Gal activity and secreted clusterin levels. sST2 is selectively secreted by decidual cells. (**C**) Impact of BCM^−^ or BCM^−^ pre-incubated with HYAL2 on uNK cell-dependent killing of senescent decidual cells in 4 biological repeat experiments. The data show relative change in the indicated measurements compared to decidual-uNK cell co-cultures exposed to UCM. Different letters above the bar graphs (mean ±SEM) indicate statistical differences (*P* < 0.05) between groups (one-way ANOVA and Dunnett’s multiple comparison test). (**D**) HYAL2 immuno-reactivity in unhatched and hatched human blastocysts (left and right panels, respectively). Scale = 100 μM. (**E**) secreted HYAL2 levels in conditioned media from embryos graded morphologically as good (grade ≥BB) or poor (<BB). (**F**) Secreted HYAL2 levels in culture medium conditioned by embryos that either resulted in a positive or negative pregnancy test following single-embryo transfer. ** indicates *P* < 0.01 (unpaired *t*-test).

Expression of HYAL2 in human blastocysts was confirmed by immunocytochemistry (Fig. 4D). We also collected BCM from 43 human IVF embryos transferred on day 5 and quantified HYAL2 levels by ELISA in individual droplets (Table S2). As shown in Figure 4E, human embryos deemed of low-quality based on morphological criteria,^40^ secreted lower levels of HYAL2 when compared to good-quality (grade: ≥BB) embryos (*P* = 0.003, unpaired *t*-test). Similarly, embryos that implanted successfully following transfer secreted higher levels of HYAL2 when compared to their unsuccessful counterparts (*P* = 0.042, unpaired *t*-test; Fig. 4F).

Taken together, our results implicate uNK cells as pivotal biosensors of embryo quality and identified the level of embryo-derived HYAL2 secretion as a major determinant for successful implantation. Figure 5 summarises the proposed mechanism of embryo-dependent modulation of uNK cell activity in endometrial fate decisions at implantation.

**Figure 5.**
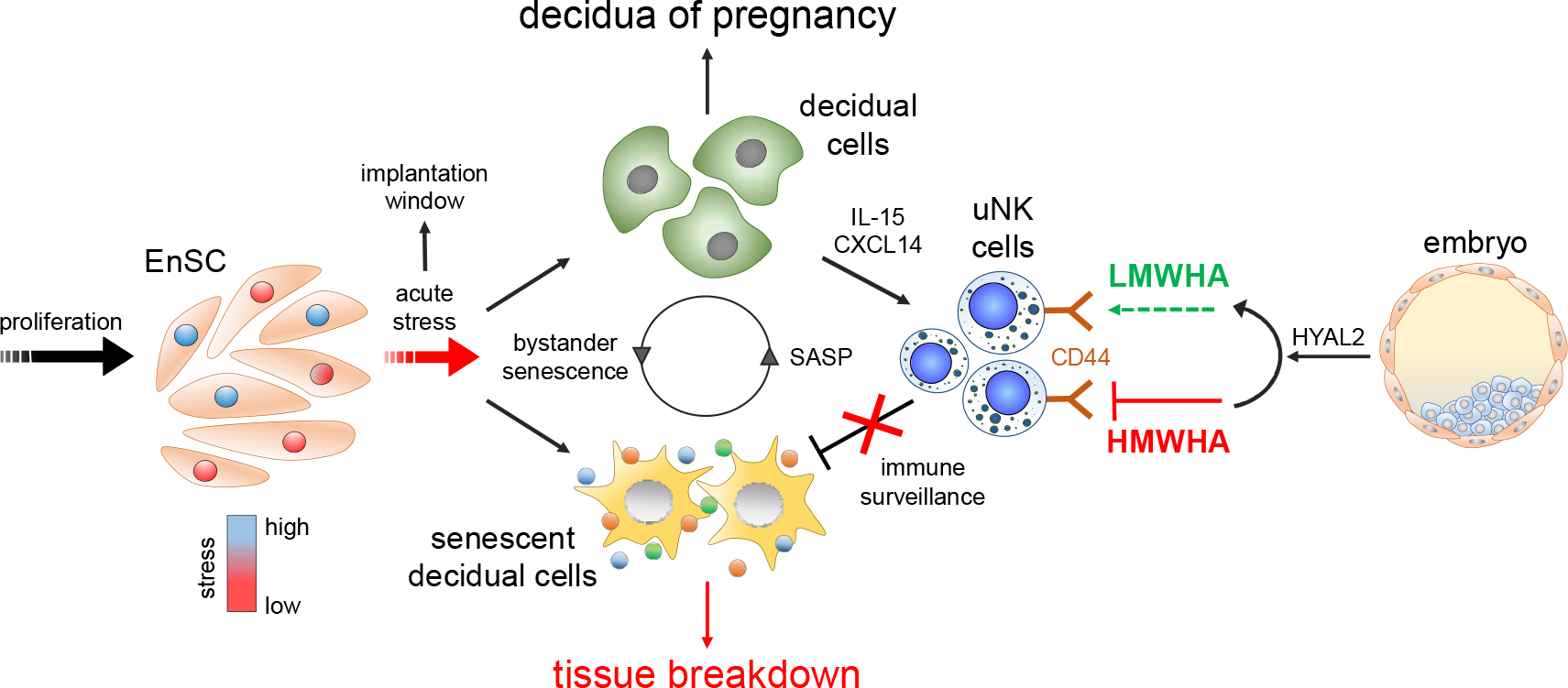
Schematic summary of the role of uNK cell sensing of embryos in endometrial fate decisions at implantation. For detailed explanation, see text.

## Discussion

Based on sensitive urinary hCG measurements, several prospective observational studies reported miscarriage rates of approximately 30% in young women trying to conceive.^41–44^ Most of these losses involve menstruation-like shedding of the endometrium one to two weeks after implantation and are therefore often not recognised clinically. The high attrition rate in the very early stages of pregnancy is widely attributed to embryonic aneuploidy, caused either by age-dependent meiotic errors in oocytes or age-independent erroneous mitotic divisions during pre-implantation development.^45,46^ A recent analysis of published single-cell RNA-seq data concluded that 80% of human embryos harbour aneuploid cells.^47^ This intrinsic genetic instability of human embryos, which gives rise to a vast array of chromosomal errors, poses a major challenge to the endometrium; how to eliminate embryos of low-fitness without compromising implantation of high-quality embryos?

Evidence from multiple mammalian species indicates that the endometrium is capable of mounting an implantation response tailored to the developmental potential of individual embryos.^21,48–50^ In humans, directed migration of decidualizing EnSC and active encapsulation of the conceptus constitutes an important initial step in the implantation process.^49^ *In vitro* studies demonstrated that high-quality human embryos stimulate migration of decidual cells whereas low-quality embryos fail to do so. Conversely, migration of undifferentiated EnSC is actively inhibited by high-quality embryos, but not low-quality embryos.^3,4,20,51^ Further, cross-species experiments demonstrated that flushing of the mouse uterus with spent culture medium from human IVF embryos deemed of insufficient quality for transfer triggers an acute cellular stress response; whereas uterine flushing with spent medium of successfully implanted human embryos (i.e. resulting in ongoing pregnancies) enhances endometrial expression of multiple metabolic and implantation genes.^21^ These observations indicate that human embryos are subjected to both negative and positive selection at implantation.^22^ The nature of embryonic ‘quality’ signals decoded by decidual cells is largely unknown, although recent studies have provided some insights. For example, compelling experimental evidence implicated embryo-derived microRNAs, more specifically hsa-miR-320a, in stimulating directional migration of decidualizing EnSC.^20^ Interestingly, hsa-miR-320a also acts in an autocrine manner to promote development of pre-implantation embryos.^52^ On the other hand, heightened secretion of tryptic serine proteinases by low-quality embryos, perhaps reflecting proteotoxic stress induced by aneuploidy, induces endoplasmic reticulum stress in decidualizing EnSC,^21^ with loss of secretion of pro-implantation modulators such as LIF, IL-1β, and HB-EGF.^14^

Despite these emerging insights, it remains unclear how a local embryo-induced endometrial stress response triggers a concatenation of events that ultimately leads to breakdown of the entire superficial endometrium, even in women receiving progesterone supplements. Several lines of evidence indicate that endometrial fate decisions at implantation pivot on the surveillance and clearance of senescent decidual cells by uNK cells. First, senescent decidual cells are progesterone-resistant and produce a cocktail of factors involved in ECM remodelling, including multiple proteases, and inflammation.^15^ Second, unlike undifferentiated EnSC, decidual cells are highly susceptible to secondary senescence (also termed ‘bystander’ senescence), enabling rapid propagation of the senescent phenotype throughout the tissue.^8,15^ We further demonstrated that recurrent pregnancy loss is associated with a pro-senescence decidual response in non-conception cycles, linked to loss of clonogenic progenitor cells and uNK cell deficiency.^15,53–55^ Here we show that uNK cell-dependent clearance of senescent decidual cells is actively suppressed by low-fitness embryos, which is predicted to promulgate secondary senescence and lead to tissue breakdown. We also demonstrate that HYAL2 restores killing of senescent decidual cells in the presence of cues from low-quality embryos. Combined with other lines of evidence presented in this study, this observation firmly implicate HMWHA-silencing of CD44 activity in abrogating uNK cell cytotoxicity. Indeed, HMWHA possesses multivalent sites to bind CD44, promotes clustering of the receptor, and inhibits downstream pathways, including Rac-dependent signalling.^24^ Interestingly, both HA production and HYAL2 expression have been shown to be critical for embryo development in animal studies.^25^ For example, inhibition of HA synthesis by 4-methyumbelliferone suppresses bovine blastocyst formation,^56^ whereas HA supplementation of culture media improves embryo development in multiple species.^25^ The effects of HA on embryo development are dependent on HYAL2 as LMWHA promotes development whereas HMWHA exerts the opposite effect.^57^ *HYAL2* transcripts are constitutively expressed in human preimplantation embryos.^26^ Here, we showed by immunocytochemistry that this glycosylphosphatidylinositol-anchored enzyme localises to both the trophectoderm and inner cell mass of human blastocysts. Importantly, HYAL2 was also detectable in BCM, with levels correlating to both embryo morphology and implantation competence. Thus, as reported for hsa-miR-320a,^20^ embryonic HYAL2 expression appears critical for both preimplantation development and endometrial responses upon implantation.

Increased deposition of HA into the ECM, specifically LMWHA, is a hallmark of inflammatory processes.^24^ In the postovulatory endometrium, HA deposition first peaks during the midluteal window of implantation but then declines rapidly in the stroma.^58^ These observations are congruent with the decline in HA production in EnSC decidualized *in vitro* for 4 days or longer. Further, loss of HA expression in decidual cells surrounding the conceptus has also been reported in mice.^59^ Although we demonstrated that HA production declines in parallel with transcriptional silencing of *HAS2*, decidualization is also associated with loss of O-linked N-acetylglucosamine (GlcNAc) transferase (OGT),^13^ an enzyme that strongly increases HAS2 activity and stability through O-GlcNAcylation of serine 221.^60^ Despite the lack of HA, uNK cell-dependent killing of senescent decidual cells was attenuated upon CD44 inhibition and addition of LMWHA did not enhance uNK cell cytotoxicity in co-cultures. These observations suggest that decidual cells either continue to produce low levels of HA fragments or express alternative CD44 ligands. Two plausible candidates are osteopontin and serglycin,^61^ both of which are strongly upregulated upon decidualization, at least at transcript level.^15^

In summary, we report for the first time that uNK cells are biosensors of low-fitness human embryos. We demonstrated that the embryo-sensing function of uNK cells relates to HYAL2 expression in the blastocyst and its ability to degrade HMWHA. Further, the role of uNK cells in embryo selection at implantation may depend on an adequate decidual response, creating a permissive local environment devoid of HMWHA production. We posit that loss of uNK cell activity at the implantation site propagates decidual senescence and facilitates menstruation-like shedding of the endometrium.

## Experimental procedures

### Endometrial sample collection

The collection of endometrial biopsies for research purposes was approved by the NHS National Research Ethics – Hammersmith and Queen Charlotte’s & Chelsea Research Ethics Committee (REC reference: 1997/5065) and Tommy’s National Reproductive Health Biobank (REC reference: 18/WA/0356). Endometrial biopsies were obtained using a Wallach Endocell^®^ endometrial sampler following transvaginal ultrasonography from patients attending a dedicated research clinic at University Hospitals Coventry and Warwickshire (UHCW) National Health Service (NHS) Trust, Coventry, UK. Written informed consent was obtained prior to tissue collection in accordance with the guidelines of the Declaration of Helsinki, 2000. Endometrial biopsies were timed 5 to 10 days after the pre-ovulatory LH surge. A total of 93 endometrial biopsies were processed for EnSC cultures, uNK cell isolation, or both. Demographic details of the tissue samples used in different experiments are shown in Table S1.

### Primary EnSC cultures

Human EnSC were isolated from endometrial biopsies as described previously.^62^ Briefly, endometrial biopsies were subjected to enzymatic digestion using 500□μg/mL collagenase type Ia (Sigma-Aldrich, Poole, UK) and 100□ μg. mL DNase I (Lorne Laboratories Ltd, Reading, UK) for 1 hr at 37°C. Digested tissue was filtered through a 40□μM cell strainer to remove glandular cell clumps, and the flow-through collected and cultured in DMEM/F12 (Thermo Scientific, Loughborough, UK) containing 10% dextran-coated charcoal-treated foetal bovine serum (DCC-FBS), 1 × antibiotic-antimycotic mix, 10 μM L-glutamine (Thermo Scientific), 1 nM estradiol and 2 μg/mL insulin (Sigma-Aldrich). Cells were lifted with 0.05% trypsin and re-seeded as required. To induce decidual transformation, confluent EnSC monolayers were down-regulated in phenol-free DMEM/F□12 media containing 2% DCC□FBS and decidualized with 10 μM medroxyprogesterone acetate (MPA) and 0.5 mM 8 bromo cAMP (C+M treatment) (Sigma Aldrich). To study the effect of exogenous HA, decidualizing EnSC were treated with either 100 μg. ml, HMWHA (molecular mass ~1320 kDa) or LMWHA (molecular mass ~33.0 kDa), purchased from Bio-Techne (USA), as described elsewhere.^63,64^ Medium was refreshed every 2 days. All experiments were performed at passage 2.

### Isolation and culture of uterine natural killer cells

uNK cells were isolated and cultured as described in detail elsewhere.^8,15^ Briefly, unattached cells from freshly digested endometrial biopsies were collected following overnight culture, and red blood cells depleted via Ficoll-Paque density medium centrifugation (400 × g) for 30 min at room temperature (RT). uNK cells were isolated through magnetic activated cell sorting (MACS) using microbeads targeting CD56 (Miltenyi Biotec, Bergisch Gladbach, Germany, 130-050-401), a specific marker for uNK cells. Cell density was adjusted to a maximum of 1×10^6^ cells per 100 μL of wash buffer (0.5% BSA, 2 mM EDTA, PBS) and incubated for 15 min at 4°C with CD56 microbeads (1:20 dilution). Unbound microbeads were removed by addition of 1 mL wash buffer, and cells were centrifuged (275 × g) for 5 min at RT and then resuspended in 500 μL wash buffer. Cells were fed through MS columns within a magnetic field wherein CD56^+^ cells were retained in columns and the negative fraction passed through. The MS columns were rinsed 3-times with 500 μL wash buffer within the magnetic field before the CD56^+^ cells were extracted by flushing the column with 1 mL wash buffer away from the magnetic field. Isolated uNK cells were cultured in DMEM/F12 media supplemented with 10% DCC-FBS, 1% L-glutamine and 1% antibiotic/antimycotic, and containing 125 pg/mL recombinant IL-15 (Sigma-Aldrich) to aid uNK cell maturation. uNK cells were cultured for no longer than 4 days.

### uNK-decidual co-culture assay

The uNK cells killing assay has been described in detail previously.^8,15^ Briefly, EnSC were seeded at a density of 50,000 cells/well in 96-well plates and decidualized for 6 days. Isolated uNK cells from whole endometrial tissue (n = 25,000) were then added to the cultures in the presence of C+M. Cultures continued for a further 2 days before harvesting for either RNA extraction or SA-β-Gal activity measurement. For experiments investigating the expression of *CLU* and *IL1RL1*, experiments were scaled to 6-well plates. Conditioned media was collected prior to lysis of cells. Where appropriate, 100 μg/mL HMWHA or LMWHA (Bio-Techne, MN, USA) or BCM were added simultaneously with uNK cells at the indicated timepoints. When indicated, HMWHA or BCM^−^ was pre-incubated with 300 IU/mL recombinant HYAL2 (Sigma-Aldrich) for 30 min prior to addition to co-cultures.

### Flow cytometry

The viability of isolated uNK cells was determined by incubation with Fixable Viability Stain 660 (FVS660, BD Biosciences, NJ, USA) diluted 1:1000 in PBS for 10 min at 4°C. For analysis of purity, cell suspensions were incubated with CD45-FITC (555482, 1:5, BD Bioscience) and CD56-PE (555516, 1:5, BD Bioscience) for 30 min in the dark at 4°C. After 2 washes with FACS buffer (0.5% BSA, 2 mM EDTA in PBS), cells were resuspended in 500 μL FACS buffer prior to analysis on BD FACSMelody™ flow cytometer.

### Immunocytochemistry of EnSC and uNK cells

Confluent EnSC monolayers were fixed with 4% paraformaldehyde for 5 min at RT and then incubated with 1% BSA in PBS-T (PBS plus 0.1% Tween 20) for 30 min at RT. Cells were stained with biotin-labelled HABP primary antibody (AMSBIO, Abington, UK, AMS.HKD-BC41, 1:125) overnight at 4°C and stained with fluorescein-conjugated streptavidin (Vector Laboratories, Peterborough, UK, 1:500). Cells were counterstained with ProLong^®^ Gold Antifade Reagent with DAPI for nuclei staining (Cell Signalling Technology, MA, USA) before image capture with an EVOS FL Auto fluorescence microscope (Life Technologies, Paisley, UK). A total of 3 images were captured for each biological repeat experiment, using matched microscope and contrast/brightness settings and quantified by fluorescence intensity using imageJ. HABP was expressed as a percentage of DAPI fluorescence to normalise for cell number in field of view.

uNK cell cytochemistry was performed with cytospin as described previously.^8^ Briefly, cell suspensions were loaded into EZ Single Cytofunnel (Thermo Scientific) and adhered to Surgipath X-tra microslides (Leica Biosystems, Wetzlar, Germany) by centrifugation (150 × g) for 5 min at RT. Cells were fixed in 4% paraformaldehyde for 10 min, washed 3-times with wash buffer (0.05% Tween 20 in PBS) and blocked (1% BSA in PBS, 30 min, RT) to prevent non-specific binding. Cells were incubated overnight at 4°C with antibodies directed against CD44 (Thermo Scientific, MA4400; 1:200) and CD56 (Leica Biosystems, NCL-L-CD56-504; 1:100). Unbound primary antibody was removed by washing and cells stained for 1 hr at RT with Alexa Fluor-488 anti-rat secondary antibody (Thermo Scientific, A11006, 1:1000) for CD44 and Alexa Fluor-594 anti-mouse secondary antibody (Thermo Scientific; A21205, 1:1000) for CD56. Cells were counterstained with ProLong^®^ Gold and viewed using an EVOS FL microscope (Thermo Scientific).

### Reverse transcription quantitative PCR (RT-qPCR)

Total RNA from human EnSC was extracted using STAT-60 (AMSBio). Recovered RNA was quantified using a Nanodrop spectrophotometer and equal amounts of total RNA were transcribed into cDNA using the QuantiTect Reverse Transcription Kit (Qiagen, Manchester, UK). Analysis of target gene expression was performed through Power SYBR Green Master Mix (Life Technologies) on a 7500 Real-Time PCR System (Applied Biosystems, CA, USA). The expression level of each gene was calculated using the ΔΔCt method and normalised against levels of the *L19* housekeeping gene. Primer sequences used were as follow: *PRL* sense 5’-AAG CTG TAG AGA TTG AGG AGC AAA C-3’, *PRL* antisense 5’-TCA GGA TGA ACC TGG CTG ACT A-3’, *IGFBP1* sense 5’-CGA AGG CTC TCC ATG TCA CCA-3’, *IGFBP1* antisense 5’-TGT CTC CTG TGC CTT GGC TAA AC-3’; *IL1RL1* sense *5’-TTG TCC TAC CAT TGA CCT CTA CAA-3’, IL1RL1* antisense *5’-GAT CCT TGA AGA GCC TGA CAA-3’, CLU* sense 5’-GGG ACC AGA CGG TCT CAG-3’, *CLU* antisense 5’-CGT ACT TAC TTC CCT GAT TGG AC-3’; *L19* sense 5’-GCG GAA GGG TAC AGC CAA-3’, *L19* antisense 5’-GCA GCC GGC GCA AA-3’; *HAS2* sense 5’-GAT GCA TTG TGA GAG GTT TCT ATG-3’, *HAS2* antisense 5’-GTA GCC AAC ATA TAT AAG CAG-3’; *HYAL2* sense 5’-CAC AGT TCC TGA GCT GGT G-3’, *HYAL2* antisense 5’-ACC AGG GCC AAT GTA ACG GT-3’.

### Enzyme-linked immunosorbent assay

Levels of secreted clusterin and sST2 in cell culture supernatant were quantified using Duoset Enzyme-linked immunosorbent assay (ELISA) kits (Biotechne, MN, USA). EnSC in 96-well plates were decidualized for 8 days, with spent media collected and refreshed every 2 days. Conditioned media was centrifuged (10000 × g) for 5 min at 4°C to remove cellular debris and stored at −20°C for assay. Quantitation of secreted factors proceeded exactly as per manufacturer’s instructions with the absorbance value determined on a PHERAstar FS plate reader (BMG Labtech, Ortenberg, Germany) at 450 nm. Samples were interpolated from known standards using a 4-parameter logistics fit.

### SA-β-Gal quantitation

Levels of cellular SA-β-Gal were determined using the 96-well Quantitative Cellular Senescence Assay kit (Cell Biolabs Inc; CA, USA), using a modified version of the manufacturer’s protocol. Decidualizing EnSC in 96-well plates were washed with ice-cold PBS and lysed in 50 μL ice-cold assay lysis buffer containing protease inhibitors (cOmplete Protease Inhibitor Cocktail, Roche). Lysates (25 μL) were transferred to black-walled, black-bottomed 96-well plates and 25 μL 2 × assay buffer added. Plates were sealed and the reaction incubated for 1 h at 37°C in a non-humidified, non-CO_2_ incubator. The reaction was terminated by the addition of 200 μL stop solution and fluorescent intensity unit (FIU) determined on a PHERAstar FS plate reader (BMG Labtech) at 360/465 nm. Assays were normalized to seeding density.

### Collection of blastocyst conditioned media

Blastocyst conditioned media (BCM) from IVF embryos cultured in individual droplets were obtained from two different units: Centre for Reproductive Medicine, UHCW NHS Trust, and Eugin Clinic, Barcelona, Spain. Following embryo transfer, BCM obtained from embryos that resulted in a positive or negative pregnancy test were designated positive (BCM^+^) or negative (BCM^−^), respectively. Parallel droplets without embryos were also obtained and designated unconditioned media (UCM).

Collection of BCM at UHCW NHS Trust was approved by the Research Ethics Committee West Midlands, Coventry and Warwickshire (REC Reference: 12/WM/0094). Normally fertilised zygotes were cultured in Cleav™ medium (Origio, Denmark) for 3 days, followed by Blast™ medium (Origio) for culture to the blastocyst stage. Cultures were performed in an atmosphere of 6% CO_2_, 5% O_2_, 89% N_2_, in an Embryoscope time-lapse incubator, with each embryo in an individual well containing 25 μL of culture medium under mineral oil. Embryo transfer was performed on day 5 after egg collection with the medium frozen *in situ* the following day and stored at −80°C until use.

Collection of BCM at Eugin Clinic, Barcelona, was approved by the Comite de Etica de la Investigación con Medicamentos (CEUGIN-2019-08-KILLDROP) on 22/10/2019. Embryos were cultured in SAGE 1-Step media (CooperSurgical, Denmark) covered with Ovoil (Vitrolife, Sweden). Cultures were performed in conventional incubators at 37°C under an atmosphere of 6% CO_2_, 5% O_2_, 89% N_2_. BCM (10 μL) was collected after embryo transfer on day 5 and transferred from the culture plate to a 0.6 mL Eppendorf. Drops were collected individually. In the same way, UCM were collected from the sample plate. BCM and UCM were stored at −80°C until use.

At both centres, the collection of BCM was fully anonymised in accordance with the approvals of the respective Ethics Committees.

### HYAL2 expression and secretion in human embryos

The research use of human cryopreserved embryos that were surplus to patient requirements was approved by the Research Ethics Committee, Coventry and Warwickshire (REC Reference: 04/Q2802/26), and the Human Fertilisation and Embryology Authority (Reference: R0155). All patients provided informed written consent.

To examine the expression of HYAL2, vitrified human blastocysts were warmed using the Vit Kit^®^ thawing protocol according to manufacturer’s instructions (Irvine Scientific, Co. Wicklow, Ireland). Blastocysts were transferred to PBS solution and fixed in 2% PF A, 4% BSA in PBS solution at RT for 2 h. Where appropriate, the zona pellucida was removed by brief incubation in acid Tyrode’s solution prior to fixation. Fixed blastocysts were washed in 4% BSA in PBS before permeabilization in 0.5% Triton X-100 in PBS at RT for 2 h and then washed in 0.2% Tween 20 in PBS for 15 min. Non-specific antibody binding sites were blocked with blocking solution (4% BSA, 5% goat serum, 0.2% Tween 20 in PBS) at RT for 1 h. Immuno-detection proceeded by overnight incubation of rabbit anti-HYAL2 primary antibody (Abcam, ab68608, 1:100) in blocking solution at 4°C. Unbound primary antibody was removed the following day by washing in 0.2% Tween 20 in PBS and blastocysts incubated for 2 hr at RT with goat anti-rabbit Alexa-488 conjugated secondary antibodies (Abcam, ab150077). Following washing, blastocysts were transferred to 5 μL Vectashield mounting medium containing DAPI on a glass slide. To protect the 3D integrity of the blastocyst, a small pinhead-sized drop of paraffin wax was placed in each corner of a 10×10 mm coverslips before setting over the sample. HYAL2 immunofluorescence in blastocysts was viewed on an Olympus 1X81 microscope using 490 nm excitation and 525 nm emission wavelengths. DAPI was detected at 359/461 nm.

To measure secreted HYAL2 levels, embryo conditioned media was diluted 1:10 with blast™ medium (Origio) with HYAL2 levels quantified by ELISA as per manufacturer’s protocols (ABIN841408, Antibodies Online). Additional information on these embryos is presented in Table S2. Morphological assessment of embryos was based on standardized criteria.^40^ Briefly, the inner cell mass and trophectoderm of embryos were assessed morphologically and graded separately on a scale of A to E, with A being the best and E the worst.^40^ Blastocysts graded <BB were deemed low quality.

### Statistical Analysis

Because of intrinsic variability between primary cultures, the data are presented as foldchange relative to an informative comparator and analysed using paired one-way ANOVA with a Dunnett’s multiple comparison test. Normal distribution was confirmed by the Shapiro-Wilk test of normality. The level of embryonic HYAL2 secretion between groups was compared using an unpaired *t*-test. Individual data points are shown for biological replicates and repeat experiments. *P* < 0.05 was considered significant with different letters above the bar graphs indicating significant differences between groups

## Acknowledgments

We are grateful to all the women who participated in this research and those who facilitated their participation. This work was supported by funds from the Tommy’s National Miscarriage Research Centre and Wellcome Trust Investigator Award to J.J.B (212233/Z/18/Z). G.M.H. was supported by the WPH Charitable Trust and UHCW NHS Trust. A.A.O. was supported by the Secretary for Universities and Research of the Ministry of Economy and Knowledge of the Government of Catalonia (GENCAT 2018 DI 103); G.T. was funded by the Torres Quevedo Program of the Spanish Ministry of Science and Innovation. This study was further supported by Clinica Eugin intramural funding.

## Author Contributions

Conceptualization, J.J.B.; Investigation, C.-S. K., P.J.B, A.A.O., S.T., T.T., J.M., E.S.L., E.S.; Writing – Original Draft, P.J.B., C.-S. K., and J.J.B.; Funding Acquisition, J.J.B. and G.H.; Resources, J.J.B., G.H., A.A.F.-N., G.T., and R.V.; Supervision, J.J.B, G.H., G.T., R.V. and P.J.B.

## Supplementary Information

**Figure S1.**
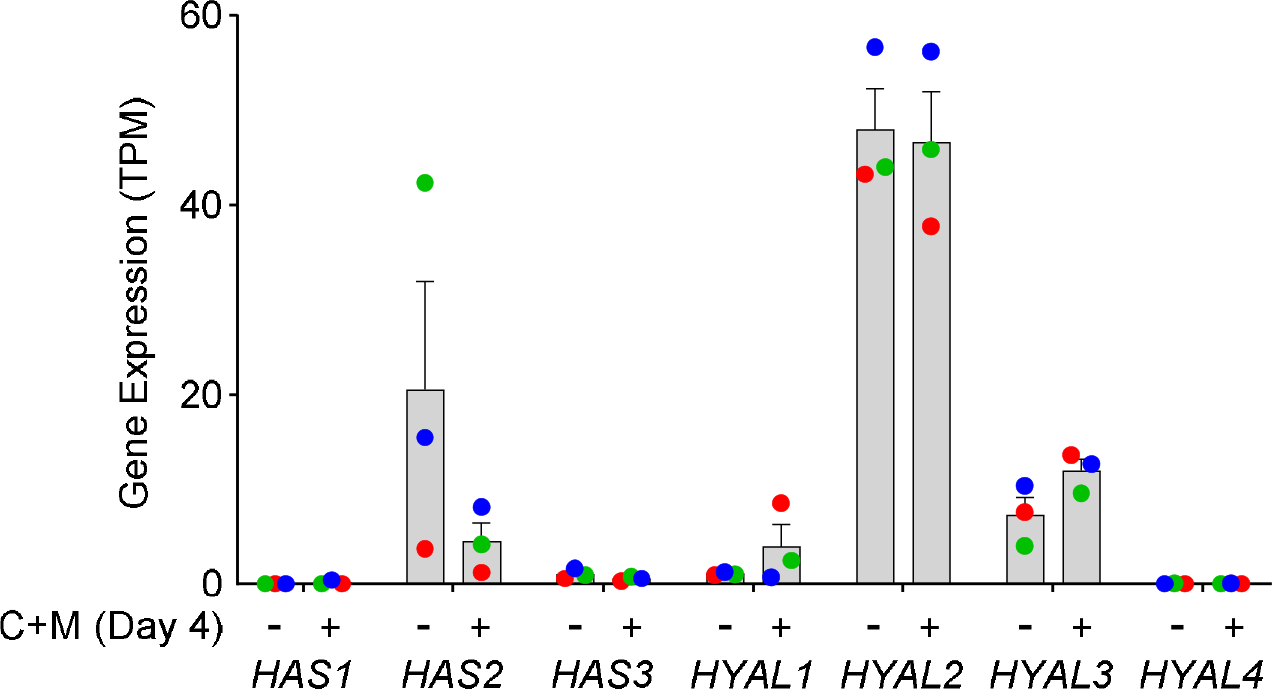
Expression of genes coding HA synthase and catabolic enzyme isoforms in undifferentiated EnSC (day 0) and cells decidualized with 8-br-cAMP and MPA (C+M) for 4 days. Individual data points from 3 biological repeat experiments are shown with bar graphs denoting mean ± SEM.

**Figure S2.**
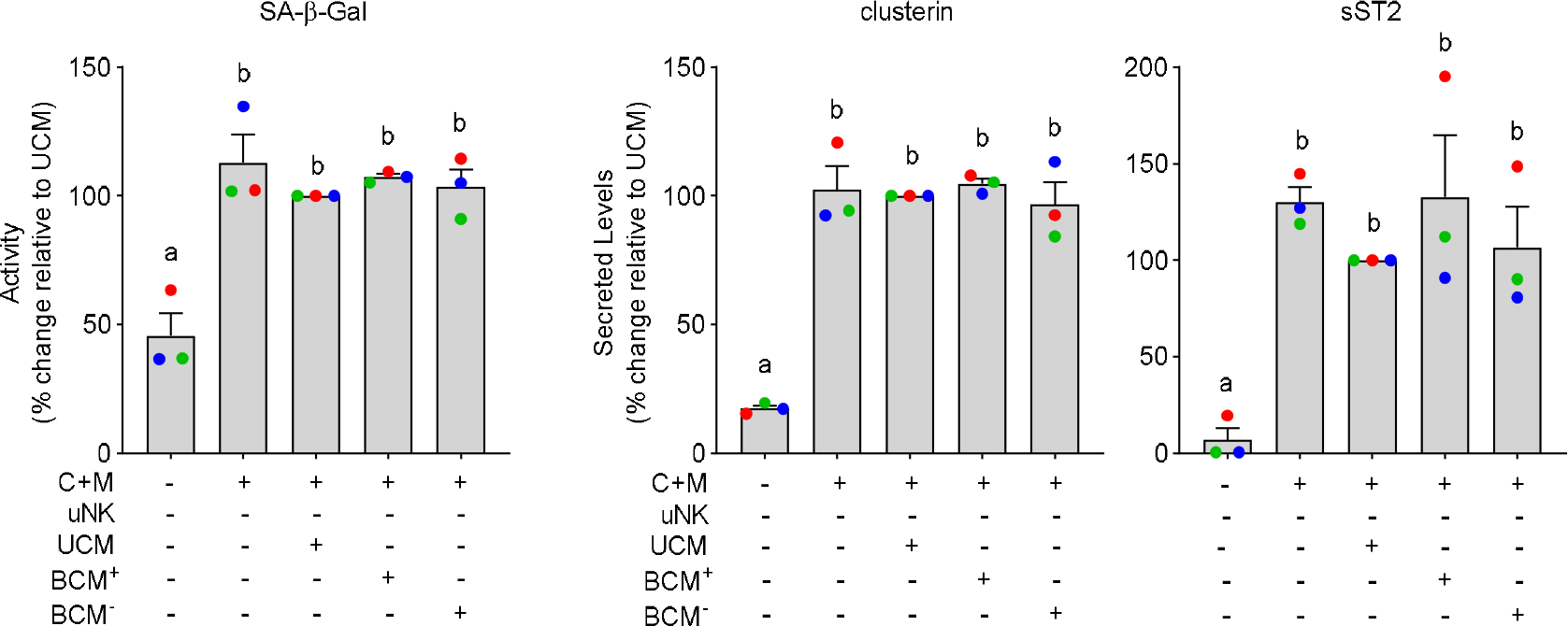
BCM does not impact on decidual subpopulations. Primary EnSC were first decidualized with C+M for 6 days and then exposed to UCM, BCM^+^ and BCM^−^, diluted 1 in 7.5 in differentiation medium (i.e. containing C+M). After 48 h, the cultures were harvested. SA-β-Gal activity and clusterin secretion mark the level of senescent decidual cells whereas sST2 secretion is confined to decidual cells. Individual data points from 3 biological repeat experiments are shown with bar graphs denoting mean ± SEM. Different letters indicate statistical differences (*P* < 0.05) between groups (one-way ANOVA and Dunnett’s multiple comparison test).

**Table S1.** Patient demographics.

| Endometrial biopsies used for EnSC cultures |  |  |  |  |
| --- | --- | --- | --- | --- |
| Figure | (n) | Age | BMI | LH+ |
| 1A-B | 4 | 31.5 (28.8-36) | 27 (24.9-29.3) | 8 (8-8.3) |
| 1C | 5 | 31 (31-32.5) | 23 (22.5-24.5) | 9 (9-10) |
| 1E-G | 3 | 38 (35.5-38) | 23 (21.5-24) | 9 (8-9.5) |
| 2G | 3 | 33 (31.5-36.5) | 23 (22-24) | 9 (8-9) |
| 2H-I | 3 | 34 (33-35.5) | 29 (29-29.8) | 7 (7-8.5) |
| 3A | 7 | 33.5 (30.8-39.8) | 24 (23-24) | 7 (7-10) |
| 3B | 5 | 35.5 (35-37.8) | 22 (21.3-22.8) | 8 (7-9) |
| 3C | 4 | 35.5 (32.8-38) | 24 (22.5-26) | 8.5 (6.75-10.5) |
| 4B | 9 | 36 (34-40) | 25 (22.5-31) | 9 (8-10) |
| 4C | 4 | 35 (33-37.5) | 26 (24.5-30.6) | 9 (8-10) |
| S2 | 3 | 36 (34-38.5) | 23.3 (22.2-25.4) | 10 (9-10.5) |
| Endometrial biopsies used for uNK cell isolation |  |  |  |  |
| 2B-C | 12 | 36 (32-37) | 23 (21.8-23.8) | 9 (8.5-9.3) |
| 2D | 3 | 43 (39-43) | 21.8 (20.9-21.9) | 7 (6.5-8.5) |
| 2E | 4 | 33 (29.5-35.8) | 25 (23.5-27) | 10 (9.3-10.3) |
| 2G | 4 | 37.5 (36.8-38.8) | 25.95 (21.8-29) | 6.8 (6.8-8.5) |
| 2I | 4 | 35.5 (34.3-37.3) | 23 (22-24) | 9 (9-9.5) |
| 3A | 9 | 36 (32-40) | 27 (24-29) | 9 (8-9) |
| 3B | 6 | 34 (31.3-39) | 27.4 (24.8-30.5) | 8.5 (7.3-9) |
| 3C | 12 | 33 (31-37.8) | 29 (24-28.8) | 9 (7-10) |
| 4B | 11 | 35 (33-38) | 24 (22.5-25.5) | 7 (6-7) |
| 4C | 11 | 32 (31.5-33.5) | 23 (22.3-23.5) | 6 (6-6.5) |

All data are median (interquartile range, Q1-Q3).

**Table S2.** Embryo grading and patient demographics (Fig. 4E-F).

|  | Good Quality (n=23) | Poor Quality (n=20) | P value |
| --- | --- | --- | --- |
| Grade | ≥BB | <BB |  |
| Age* | 33 (31-35) | 36 (31.5-39.3) | 0.314 |
| IVF | 17 | 6 |  |
| ICSI | 6 | 14 |  |

|  | Pregnant (n=20)^ | Non-Pregnant (n=22) | P value |
| --- | --- | --- | --- |
| Age* | 32.1 (30-34) | 36.3 (33-40) | 0.003 |
| IVF | 16 | 6 |  |
| ICSI | 4 | 16 |  |

\*median (interquartile range, Q1-Q3)
^ One ectopic pregnancy excluded
Significance determined by *t*-test

## References

1 Gellersen, B. & Brosens, J. J. Cyclic decidualization of the human endometrium in reproductive health and failure. Endocr Rev 35, 851–905, doi:10.1210/er.2014-1045 (2014).

2 Haig, D. Genetic conflicts in human pregnancy. Q Rev Biol 68, 495–532, doi:10.1086/418300 (1993).

3 Berkhout, R. P. et al. High-quality human preimplantation embryos actively influence endometrial stromal cell migration. J Assist Reprod Genet 35, 659–667, doi:10.1007/s10815-017-1107-z (2018).

4 Weimar, C. H. et al. Endometrial stromal cells of women with recurrent miscarriage fail to discriminate between high- and low-quality human embryos. PLoS One 7, e41424, doi:10.1371/journal.pone.0041424 (2012).

5 Erlebacher, A. Immunology of the maternal-fetal interface. Annu Rev Immunol 31, 387–411, doi:10.1146/annurev-immunol-032712-100003 (2013).

6 Erkenbrack, E. M. et al. The mammalian decidual cell evolved from a cellular stress response. PLoS Biol 16, e2005594, doi:10.1371/journal.pbio.2005594 (2018).

7 Al-Sabbagh, M. et al. NADPH oxidase-derived reactive oxygen species mediate decidualization of human endometrial stromal cells in response to cyclic AMP signaling. Endocrinology 152, 730–740, doi:10.1210/en.2010-0899 (2011).

8 Brighton, P. J. et al. Clearance of senescent decidual cells by uterine natural killer cells in cycling human endometrium. Elife 6, doi:10.7554/eLife.31274 (2017).

9 Mor, G., Cardenas, I., Abrahams, V. & Guller, S. Inflammation and pregnancy: the role of the immune system at the implantation site. Ann N Y Acad Sci 1221, 80–87, doi:10.1111/j.1749-6632.2010.05938.x (2011).

10 Salker, M. S. et al. Disordered IL-33/ST2 activation in decidualizing stromal cells prolongs uterine receptivity in women with recurrent pregnancy loss. PLoS One 7, e52252, doi:10.1371/journal.pone.0052252 (2012).

11 Brosens, J. J., Hayashi, N. & White, J. O. Progesterone receptor regulates decidual prolactin expression in differentiating human endometrial stromal cells. Endocrinology 140, 4809–4820, doi:10.1210/endo.140.10.7070 (1999).

12 Leitao, B. et al. Silencing of the JNK pathway maintains progesterone receptor activity in decidualizing human endometrial stromal cells exposed to oxidative stress signals. FASEB J 24, 1541–1551, doi:10.1096/fj.09-149153 (2010).

13 Muter, J. et al. The Glycosyltransferase EOGT Regulates Adropin Expression in Decidualizing Human Endometrium. Endocrinology 159, 994–1004, doi:10.1210/en.2017-03064 (2018).

14 Teklenburg, G. et al. Natural selection of human embryos: decidualizing endometrial stromal cells serve as sensors of embryo quality upon implantation. PLoS One 5, e10258, doi:10.1371/journal.pone.0010258 (2010).

15 Lucas, E. S. et al. Recurrent pregnancy loss is associated with a pro-senescent decidual response during the peri-implantation window. Commun Biol 3, 37, doi:10.1038/s42003-020-0763-1 (2020).

16 Ochiai, A. et al. Resveratrol inhibits decidualization by accelerating downregulation of the CRABP2-RAR pathway in differentiating human endometrial stromal cells. Cell Death Dis 10, 276, doi:10.1038/s41419-019-1511-7 (2019).

17 Critchley, H. O. D., Maybin, J. A., Armstrong, G. M. & Williams, A. R. W. Physiology of the Endometrium and Regulation of Menstruation. Physiol Rev 100, 1149–1179, doi:10.1152/physrev.00031.2019 (2020).

18 Ewington, L. J., Tewary, S. & Brosens, J. J. New insights into the mechanisms underlying recurrent pregnancy loss. J Obstet Gynaecol Res 45, 258–265, doi:10.1111/jog.13837 (2019).

19 Haig, D. Cooperation and conflict in human pregnancy. Curr Biol 29, R455–R458, doi:10.1016/j.cub.2019.04.040 (2019).

20 Berkhout, R. P. et al. High-quality human preimplantation embryos stimulate endometrial stromal cell migration via secretion of microRNA hsa-miR-320a. Hum Reprod, doi:10.1093/humrep/deaa149 (2020).

21 Brosens, J. J. et al. Uterine selection of human embryos at implantation. Sci Rep 4, 3894, doi:10.1038/srep03894 (2014).

22 Macklon, N. S. & Brosens, J. J. The human endometrium as a sensor of embryo quality. Biol Reprod 91, 98, doi:10.1095/biolreprod.114.122846 (2014).

23 Kane, N., Kelly, R., Saunders, P. T. & Critchley, H. O. Proliferation of uterine natural killer cells is induced by human chorionic gonadotropin and mediated via the mannose receptor. Endocrinology 150, 2882–2888, doi:10.1210/en.2008-1309 (2009).

24 Tavianatou, A. G. et al. Hyaluronan: molecular size-dependent signaling and biological functions in inflammation and cancer. FEBS J 286, 2883–2908, doi:10.1111/febs.14777 (2019).

25 Fouladi-Nashta, A. A., Raheem, K. A., Marei, W. F., Ghafari, F. & Hartshorne, G. M. Regulation and roles of the hyaluronan system in mammalian reproduction. Reproduction 153, R43–R58, doi:10.1530/REP-16-0240 (2017).

26 Ruane, P. T. et al. The effects of hyaluronate-containing medium on human embryo attachment to endometrial epithelial cells in vitro. Hum Reprod Open 2020, hoz033, doi:10.1093/hropen/hoz033 (2020).

27 Takahashi, H. et al. Extravillous trophoblast cell invasion is promoted by the CD44-hyaluronic acid interaction. Placenta 35, 163–170, doi:10.1016/j.placenta.2013.12.009 (2014).

28 Ripellino, J. A., Klinger, M. M., Margolis, R. U. & Margolis, R. K. The hyaluronic-acid binding region as a specific probe for the localization of hyaluronic-acid in tissue-sections – application to chick-embryo and rat-brain. Journal of Histochemistry & Cytochemistry 33, 1060–1066, doi:10.1177/33.10.4045184 (1985).

29 Weigel, P. H., Hascall, V. C. & Tammi, M. Hyaluronan synthases. Journal of Biological Chemistry 272, 13997–14000, doi:10.1074/jbc.272.22.13997 (1997).

30 Baggenstoss, B. A. et al. Hyaluronan synthase control of synthesis rate and hyaluronan product size are independent functions differentially affected by mutations in a conserved tandem B-X-7-B motif. Glycobiology 27, 154–164, doi:10.1093/glycob/cww089 (2017).

31 Martin-DeLeon, P. A. Epididymal SPAM1 and its impact on sperm function. Molecular and Cellular Endocrinology 250, 114–121, doi:10.1016/j.mce.2005.12.033 (2006).

32 Hemming, R. et al. Mouse Hyal3 encodes a 45-to 56-kDa glycoprotein whose overexpression increases hyaluronidase 1 activity in cultured cells. Glycobiology 18, 280–289, doi:10.1093/glycob/cwn006 (2008).

33 Atmuri, V. et al. Hyaluronidase 3 (HYAL3) knockout mice do not display evidence of hyaluronan accumulation. Matrix Biology 27, 653–660, doi:10.1016/j.matbio.2008.07.006 (2008).

34 Xie, D. et al. Rewirable gene regulatory networks in the preimplantation embryonic development of three mammalian species. Genome Research 20, 804–815, doi:10.1101/gr.100594.109 (2010).

35 Sague, S. L., Tato, C., Pure, E. & Hunter, C. A. The regulation and activation of CD44 by natural killer (NK) cells and its role in the production of IFN-gamma. J Interferon Cytokine Res 24, 301–309, doi:10.1089/107999004323065093 (2004).

36 Yasuo, T., Kitaya, K., Yamaguchi, T., Fushiki, S. & Honjo, H. Possible role of hematopoietic CD44/chondroitin sulfate interaction in extravasation of peripheral blood CD16(-) natural killer cells into human endometrium. J Reprod Immunol 78, 1–10, doi:10.1016/j.jri.2007.09.002 (2008).

37 Stern, R., Asari, A. A. & Sugahara, K. N. Hyaluronan fragments: An information-rich system. European Journal of Cell Biology 85, 699–715, doi:10.1016/j.ejcb.2006.05.009 (2006).

38 Ohno-Nakahara, M. et al. Induction of CD44 and MMP expression by hyaluronidase treatment of articular chondrocytes. J Biochem 135, 567–575, doi:10.1093/jb/mvh069 (2004).

39 Alves, C. S., Yakovlev, S., Medved, L. & Konstantopoulos, K. Biomolecular Characterization of CD44-Fibrin(ogen) Binding distinct molecular requirements mediate binding of standard and variant isoforms of cd44 to immobilized fibrin(ogen). Journal of Biological Chemistry 284, 1177–1189, doi:10.1074/jbc.M805144200 (2009).

40 Gardner, D. K., Lane, M., Stevens, J., Schlenker, T. & Schoolcraft, W. B. Blastocyst score affects implantation and pregnancy outcome: towards a single blastocyst transfer. Fertil Steril 73, 1155–1158, doi:10.1016/s0015-0282(00)00518-5 (2000).

41 Foo, L. et al. Peri-implantation urinary hormone monitoring distinguishes between types of first-trimester spontaneous pregnancy loss. Paediatr Perinat Epidemiol, doi:10.1111/ppe.12613 (2020).

42 Wang, X. et al. Conception, early pregnancy loss, and time to clinical pregnancy: a population-based prospective study. Fertil Steril 79, 577–584, doi:10.1016/s0015-0282(02)04694-0 (2003).

43 Wilcox, A. J. et al. Incidence of early loss of pregnancy. N Engl J Med 319, 189–194, doi:10.1056/NEJM198807283190401 (1988).

44 Zinaman, M. J., Clegg, E. D., Brown, C. C., O’Connor, J. & Selevan, S. G. Estimates of human fertility and pregnancy loss. Fertil Steril 65, 503–509 (1996).

45 McCoy, R. C. Mosaicism in Preimplantation Human Embryos: When Chromosomal Abnormalities Are the Norm. Trends Genet 33, 448–463, doi:10.1016/j.tig.2017.04.001 (2017).

46 McCoy, R. C. et al. Evidence of Selection against Complex Mitotic-Origin Aneuploidy during Preimplantation Development. PLoS Genet 11, e1005601, doi:10.1371/journal.pgen.1005601 (2015).

47 Starostik, M. R., Sosina, O. A. & McCoy, R. C. Single-cell analysis of human embryos reveals diverse patterns of aneuploidy and mosaicism. Genome Res, doi:10.1101/gr.262774.120 (2020).

48 Bauersachs, S. et al. The endometrium responds differently to cloned versus fertilized embryos. Proc Natl Acad Sci U S A 106, 5681–5686, doi:10.1073/pnas.0811841106 (2009).

49 Biase, F. H. et al. Fine-tuned adaptation of embryo-endometrium pairs at implantation revealed by transcriptome analyses in Bos taurus. PLoS Biol 17, e3000046, doi:10.1371/journal.pbio.3000046 (2019).

50 Mansouri-Attia, N. et al. Endometrium as an early sensor of in vitro embryo manipulation technologies. Proc Natl Acad Sci USA 106, 5687–5692, doi:10.1073/pnas.0812722106 (2009).

51 Weimar, C. H., Macklon, N. S., Post Uiterweer, E. D., Brosens, J. J. & Gellersen, B. The motile and invasive capacity of human endometrial stromal cells: implications for normal and impaired reproductive function. Hum Reprod Update 19, 542–557, doi:10.1093/humupd/dmt025 (2013).

52 Feng, R. et al. MiRNA-320 in the human follicular fluid is associated with embryo quality in vivo and affects mouse embryonic development in vitro. Sci Rep 5, 8689, doi:10.1038/srep08689 (2015).

53 Fukui, A., Funamizu, A., Fukuhara, R. & Shibahara, H. Expression of natural cytotoxicity receptors and cytokine production on endometrial natural killer cells in women with recurrent pregnancy loss or implantation failure, and the expression of natural cytotoxicity receptors on peripheral blood natural killer cells in pregnant women with a history of recurrent pregnancy loss. J Obstet Gynaecol Res 43, 1678-1686, doi:10.1111/jog.13448 (2017).

54 Katano, K. et al. Peripheral natural killer cell activity as a predictor of recurrent pregnancy loss: a large cohort study. Fertil Steril 100, 1629–1634, doi:10.1016/j.fertnstert.2013.07.1996 (2013).

55 Lucas, E. S. et al. Loss of Endometrial Plasticity in Recurrent Pregnancy Loss. Stem Cells 34, 346–356, doi:10.1002/stem.2222 (2016).

56 Marei, W. F., Salavati, M. & Fouladi-Nashta, A. A. Critical role of hyaluronidase-2 during preimplantation embryo development. Mol Hum Reprod 19, 590–599, doi:10.1093/molehr/gat032 (2013).

57 Marei, W. F. A. et al. Hyaluronan and hyaluronidase, which is better for embryo development? Theriogenology 86, 940–948, doi:10.1016/j.theriogenology.2016.03.017 (2016).

58 Salamonsen, L. A., Shuster, S. & Stern, R. Distribution of hyaluronan in human endometrium across the menstrual cycle. Implications for implantation and menstruation. Cell Tissue Res 306, 335–340, doi:10.1007/s004410100452 (2001).

59 Brown, J. J. & Papaioannou, V. E. Distribution of hyaluronan in the mouse endometrium during the periimplantation period of pregnancy. Differentiation 52, 61–68, doi:10.1111/j.1432-0436.1992.tb00500.x (1992).

60 Vigetti, D. et al. Role of UDP-N-acetylglucosamine (GlcNAc) and O-GlcNAcylation of hyaluronan synthase 2 in the control of chondroitin sulfate and hyaluronan synthesis. J Biol Chem 287, 35544–35555, doi:10.1074/jbc.M112.402347 (2012).

61 Chen, C., Zhao, S., Karnad, A. & Freeman, J. W. The biology and role of CD44 in cancer progression: therapeutic implications. J Hematol Oncol 11, 64, doi:10.1186/s13045-018-0605-5 (2018).

62 Barros, F. S. V., Brosens, J. J. & Brighton, P. J. Isolation and Primary Culture of Various Cell Types from Whole Human Endometrial Biopsies. bio-protocol 6, doi:DOI: 10.21769/BioProtoc.2028 (2016).

63 Rowley, J. E. et al. Low Molecular Weight Hyaluronan Induces an Inflammatory Response in Ovarian Stromal Cells and Impairs Gamete Development In Vitro. Int J Mol Sci 21, doi:10.3390/ijms21031036 (2020).

64 Zhu, R. et al. Hyaluronan up-regulates growth and invasion of trophoblasts in an autocrine manner via PI3K/AKT and MAPK/ERK1/2 pathways in early human pregnancy. Placenta 34, 784–791, doi:10.1016/j.placenta.2013.05.009 (2013).

